# Vaxign-DL: A Deep Learning-based Method for Vaccine Design and its Evaluation

**DOI:** 10.1101/2023.11.29.569096

**Authors:** Yuhan Zhang, Anthony Huffman, Justin Johnson, Yongqun He

**Author notes:** Co-first author (Yuhan Zhang, Anthony Huffman).

## Abstract

Reverse vaccinology (RV) provides a systematic approach to identifying potential vaccine candidates based on protein sequences. The integration of machine learning (ML) into this process has greatly enhanced our ability to predict viable vaccine candidates from these sequences. We have previously developed a Vaxign-ML program based on the eXtreme Gradient Boosting (XGBoost). In this study, we further extend our work to develop a Vaxign-DL program based on deep learning techniques. Deep neural networks assemble non-linear models and learn multilevel abstraction of data using hierarchically structured layers, offering a data-driven approach in computational design models. Vaxign-DL uses a three-layer fully connected neural network model. Using the same bacterial vaccine candidate training data as used in Vaxign-ML development, Vaxign-DL was able to achieve an Area Under the Receiver Operating Characteristic of 0.94, specificity of 0.99, sensitivity of 0.74, and accuracy of 0.96. Using the Leave-One-Pathogen-Out Validation (LOPOV) method, Vaxign-DL was able to predict vaccine candidates for 10 pathogens. Our benchmark study shows that Vaxign-DL achieved comparable results with Vaxign-ML in most cases, and our method outperforms Vaxi-DL in the accurate prediction of bacterial protective antigens.

## Introduction

Vaccines are vital in preventing infectious diseases by stimulating the immune system. All infectious diseases are caused by a microbial organism known as a pathogen. A vaccine works by presenting the immune system with a protein that is used to recognize a specific infectious disease.

The chosen protein, known as a protective antigen, is typically produced by the organism responsible for the infectious disease. However, some vaccines use protective antigens that have been modified from the endogenous protein to improve the safety and efficacy of a vaccine (1-3). As such, developing an effective vaccine is a complex process that traditionally takes years of testing and trials. The emergence of high throughput sequencing and improved computing programs have enabled the field of bioinformatics to develop techniques to pre-screen candidate protective antigens. The use of the *in silico* analysis of protective candidates is known as Reverse Vaccinology (RV) (4). The successful use of RV speeds this process, with the most successful demonstration being the rapid development of the COVID-19 vaccine (5).

Reverse Vaccinology approaches can be classified into two categories: filtering-based methods such as NERVE (6) and Vaxign (7), and machine learning-based methods such as Vaxign-ML (8) and Vaxi-DL (9). Filtering methods simply assess a given protective antigen on a selected set of specific criteria to recommend if it is a good vaccine antigen target or not. The original Vaxign tool, for example, would filter proteins out if a high number of transmembrane helices were found (7). A limitation of filtering is that the algorithm assesses each criterion separately instead of holistically, causing potential misclassifications in the data. Machine learning, in contrast, is capable of looking at the same set of inputs and providing a more comprehensive prediction. The use of these algorithms in neural networks enables the possibility of improved performance compared to other conventional RV tools (7). However, the output of a machine learning algorithm is dependent on the quality of datasets provided in the study of Vaxign-ML, the successor to Vaxign, which utilizes an XGBoost method that emerged as the top-performing model (8). Vaxign-ML utilized a gold standard set of experimentally verified protective antigens (10).

Vaxi-DL (9) is a web-based deep learning server that predicts potential vaccine candidates using a fully connected neural network. Deep learning can leverage larger datasets to identify patterns that are not obvious in smaller datasets. Previously, our lab had assumed that the current set of known vaccines was too small to be sufficient to predict protective candidates. However, the Vaxi-DL paper does not include negative samples in their benchmark dataset. Our prototype data testing showed Vaxi-DL indeed generated low specificity records for bacteria samples. As such, our lab wanted to assess if the deep learning methodologies were viable for vaccine design and to iterate improvement on the dataset.

In this study, we report our development of Vaxign-DL, a new RV program based on deep learning. Vaxign-DL was developed using the same gold-standard training and testing data as recorded in the Vaxign-ML study (8). We also compared the performance of Vaxign-ML prediction model with other vaccine design programs including Vaxign (7), Vaxi-DL, VaxiJen (11), etc.

## Methods

### Overall Vaxign-DL pipeline

The overall pipeline is described in Figure 1. Positive and negative classes were assigned from the dataset previously generated for Vaxign-ML. These sequences were then retrieved and processed from Protegen (10) and UniProt databases. The biological and physicochemical attributes of these protein sequences were annotated utilizing publicly accessible bioinformatics software. Because of the inherent class imbalance in the dataset, we adopted multiple sampling strategies. The performance of these strategies was then compared to determine the optimal data handling approach. Stratified 5-fold cross-validation was implemented to assess the deep learning algorithms, coupled with feature selection and hyperparameter optimization. The model employed in this study is a Multi-layer Perceptron (MLP). This model is a category of deep learning algorithms, which operate through the sequential layering of nonlinear processing units. This mechanism allows the capture and modeling of highly nonlinear data relationships.

**Figure 1.**
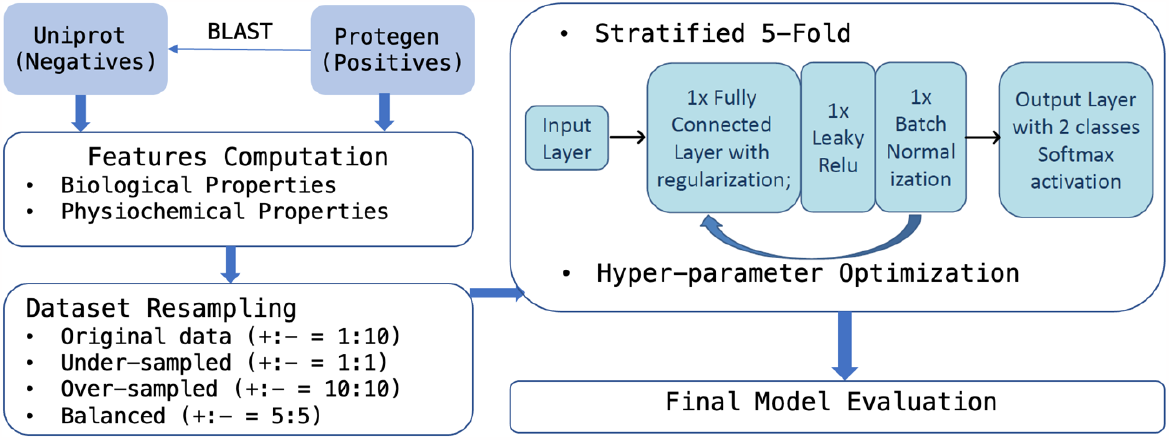
Schematic Representation of the Workflow.

### Dataset

The dataset employed for this study utilized the same feature generation techniques for Vaxign-ML (8). Specifically, the dataset used comprises 397 instances of positive samples and 3,970 instances of negative samples (+:− = 1:10), with 509 distinct features. Features include both biological and physicochemical features. We have merged Gram+ and Gram-bacterial data by adding a column signifying if it is a Gram+ or Gram-bacteria. The majority of these features correspond to physical properties of a given sequence, namely predictions of the hydrophobicity, average flexibility, polarizability, mutability, free energy, residue volume, steric pressure, and solvent accessibility of the molecule. The remaining major features are biological predictions and include localization probability of 6 possible extra-cellular locations, prediction of trans-membrane helices using TMHMM (12), and adhesin probability (13).

### Balance dataset with resampling

The initial dataset presented an imbalanced distribution that skewed towards negative class, with a positive-to-negative class ratio of 1:10. This imbalance could pose challenges for deep learning models, as they might get biased towards the majority class, leading to suboptimal predictions for the minority class. To investigate the impact of class imbalance on deep learning models and the efficacy of various resampling methods, we employed three distinct resampling techniques. Firstly, we used the under-sampling technique, where instances from the majority (negative) class were randomly discarded to equalize its count with that of the minority (positive) class, resulting in a balanced ratio of 1:1. Secondly, we incorporated the Synthetic Minority Oversampling Technique (SMOTE) (14), resulting in a ratio 10:10. This method was applied exclusively to the training dataset and not the validation or testing data, to maintain the integrity of our evaluation phases. SMOTE generates synthetic samples for the minority class, in this case, the positive class, through an interpolation process between neighboring instances. Thirdly, we adopted a balanced approach that combined the principles of both under-sampling and over-sampling by first downsampling the majority class and then synthesizing the minority class, resulting in a ratio 5:5.

### Model Construction

Our Multilayer Perceptron (MLP) model, illustrated in Figure 2, features a layered structure starting with an input layer, followed by three fully connected (FC) hidden layers. The input features were scaled using a MinMax scaler. The first two hidden layers have 256 hidden units each, and the third hidden layer has 128 hidden units. Each FC layer uses the LeakyReLU activation function with a slight slope of 0.005 to maintain activity when inputs are negative. Following each LeakyReLU function, batch normalization is applied to improve optimization and stability. To increase the robustness and prevent overfitting, dropout layers with a dropout rate of 0.1 are implemented. The final output layer consists of two nodes to identify ‘protective’ or ‘non-protective’ candidates. A softmax activation function is used at this layer to guarantee outputs that represent the probabilities for these classes, which sum up to one.

**Figure 2.**
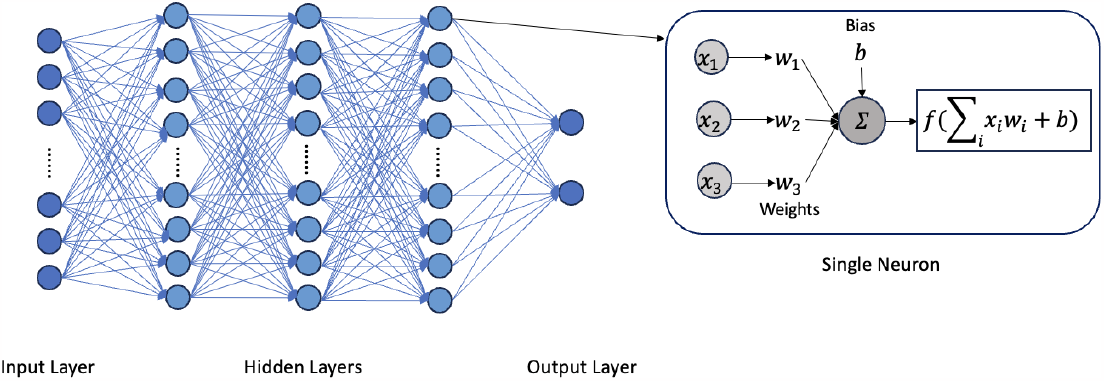
MLP model visualization

Weights are initialized using the Glorot uniform initializer. The model is trained using the Adam optimizer with learning rate 1e-3 and weight decay 1e-4 for L2 regularization. A learning rate scheduler is implemented, which adaptively adjusts the learning rate as the training progresses. Cross-entropy loss, enhanced by a label smoothing factor of 0.2, is the loss function. Early stopping is employed as a form of regularization to prevent overfitting during the training process.

### Stratified 5-fold cross-validation

The training dataset is partitioned into 5 distinct subsets while maintaining the proportional representation of both positive and negative samples. After the partitioning process, the Synthetic Minority Oversampling Technique (SMOTE) is implemented on the training dataset, ensuring that the synthetic samples do not leak into the validation set. Among the 5 subsets, 4 parts are dedicated to the training process where the model learns and adjusts its parameters, while the remaining part is reserved exclusively for the validation phase.

### Leave-one-pathogen-out validation

To provide a more general estimation of the classification performance and to simulate a scenario in which vaccine candidates would be required for a novel pathogen, we utilized the Leave-One-Pathogen-Out Validation (LOPOV) method (8). This approach involved testing a set of ten pathogens, comprising four Gram-positive pathogens (*Mycobacterium tuberculosis, Staphylococcus aureus, Streptococcus pneumoniae*, and *Streptococcus pyogenes*) and six Gram-negative pathogens (*Helicobacter pylori, Neisseria meningitidis, Brucella spp*., *Escherichia coli, Yersinia pestis*, and *Hemophilus influenzae*). For each LOPOV process, the positive and negative samples from one of these ten pathogens were exclusively held out for testing, and the remaining samples from the original dataset were employed for training the model.

### Performance evaluation

Performance evaluation was done by measuring the area under the Receiver Operating Characteristic (AUROC) and Precision-Recall Curve (AUPRC), the weighted F1 score, and the Matthews correlation coefficient (MCC) of our dataset. The latter two metrics were used to ensure that we had good results for all four confusion matrix categories (true positives, false negatives, true negatives, and false positives).

### Benchmark with an independent dataset

We utilized our gold-standard benchmark dataset from the original Vaxign-ML paper (8). More specifically, this dataset includes 131 positive protective antigens (59 Gram+ antigens and 72 Gram-antigens) and 115 negative antigens (40 Gram+ antigens and 75 Gram-antigens). These proteins were consolidated from Dalsass et al., 2019 (15) and Heinson et al., 2017 (16). This benchmark dataset removed any duplicates or proteins that were orthologs or a subset of a protein within our training dataset. In our benchmark evaluation, we merged the Gram+ and Gram-datasets together by adding a new feature called Gram staining. The GitHub link to our gold-standard benchmark data is here: https://github.com/VIOLINet/Vaxign-ML-docker/tree/master/Benchmark.

## Results

### Performance analysis of Vaxign-DL model

We initially trained Vaxign-DL on the original dataset before evaluating the effect of resampling methods on Vaxign-DL performance (Table 1). The model achieved high validation accuracy (Val accuracy) (0.96 ± 0.01) and specificity (0.99 ± 0.004); the sensitivity is comparatively low (0.74 ± 0.07). To improve sensitivity, we first over-sampled the dataset, and the average sensitivity increased marginally to (0.78 ± 0.035). Other metrics, including validation accuracy, specificity, and AUROC remain relatively unchanged (Table 1, Figure 3). Then, we under-sampled the dataset. This showed a more notable improvement in sensitivity (0.86 ± 0.053) and provided a slight increase in AUROC to 0.95 (Figure 3). However, there was a decrease in all other performance metrics, but other metrics, especially MCC (0.65 ± 0.028), declined. Lastly, we tried a balanced sampling method. This further improved sensitivity (0.87 ± 0.053), but other metrics remained low. The usage of the original dataset achieved the highest accuracy, specificity, weighted F1, MCC, and AUPRC. As such, the final model utilized the original dataset.

**Table 1.** Performance of Vaxign-DL using Vaxign-ML training data.

|  | Fold # | Val Accuracy | Sensitivity | Specificity | Weighted F1 | MCC | AUPRC | AUROC |
| --- | --- | --- | --- | --- | --- | --- | --- | --- |
| Original data | Fold 1 | 0.96 | 0.74 | 0.98 | 0.96 | 0.76 | 0.85 | 0.94 |
|  | Fold 2 | 0.95 | 0.64 | 0.99 | 0.96 | 0.74 | 0.79 |  |
|  | Fold 3 | 0.96 | 0.77 | 0.99 | 0.96 | 0.79 | 0.88 |  |
|  | Fold 4 | 0.98 | 0.85 | 0.99 | 0.98 | 0.87 | 0.91 |  |
|  | Fold 5 | 0.96 | 0.70 | 0.99 | 0.96 | 0.77 | 0.79 |  |
|  | Avg | **0.96 ± 0.008** | 0.74 ± 0.071 | **0.99 ± 0.004** | **0.96 ± 0.008** | **0.78 ± 0.046** | **0.84 ± 0.049** |  |
| Oversampled | Fold 1 | 0.96 | 0.83 | 0.98 | 0.96 | 0.78 | 0.82 | 0.94 |
|  | Fold 2 | 0.95 | 0.74 | 0.98 | 0.96 | 0.74 | 0.78 |  |
|  | Fold 3 | 0.96 | 0.77 | 0.98 | 0.96 | 0.78 | 0.86 |  |
|  | Fold 4 | 0.96 | 0.82 | 0.98 | 0.97 | 0.82 | 0.90 |  |
|  | Fold 5 | 0.96 | 0.76 | 0.97 | 0.95 | 0.73 | 0.79 |  |
|  | Avg | **0.96 ± 0.004** | 0.78 ± 0.035 | 0.98 ± 0.004 | **0.96 ± 0.005** | 0.77 ± 0.030 | 0.83 ± 0.045 |  |
| Undersampled | Fold 1 | 0.92 | 0.89 | 0.93 | 0.93 | 0.68 | 0.79 | 0.95 |
|  | Fold 2 | 0.93 | 0.71 | 0.95 | 0.93 | 0.63 | 0.74 |  |
|  | Fold 3 | 0.93 | 0.86 | 0.94 | 0.94 | 0.69 | 0.86 |  |
|  | Fold 4 | 0.94 | 0.92 | 0.95 | 0.95 | 0.75 | 0.85 |  |
|  | Fold 5 | 0.93 | 0.76 | 0.95 | 0.94 | 0.65 | 0.71 |  |
|  | Avg | 0.91 ± 0.009 | 0.86 ± 0.053 | 0.92 ± 0.011 | 0.93 ± 0.007 | 0.65 ± 0.028 | 0.80 ± 0.063 |  |
| Balanced | Fold 1 | 0.90 | 0.90 | 0.91 | 0.92 | 0.64 | 0.77 | 0.95 |
|  | Fold 2 | 0.92 | 0.83 | 0.93 | 0.93 | 0.65 | 0.79 |  |
|  | Fold 3 | 0.90 | 0.90 | 0.91 | 0.92 | 0.63 | 0.86 |  |
|  | Fold 4 | 0.92 | 0.92 | 0.93 | 0.93 | 0.69 | 0.84 |  |
|  | Fold 5 | 0.90 | 0.78 | 0.91 | 0.91 | 0.57 | 0.76 |  |
|  | Avg | 0.91 ± 0.010 | **0.87 ± 0.053** | 0.92 ± 0.010 | 0.92 ± 0.008 | 0.64 ± 0.038 | 0.80 ± 0.039 |  |

**Figure 3.**
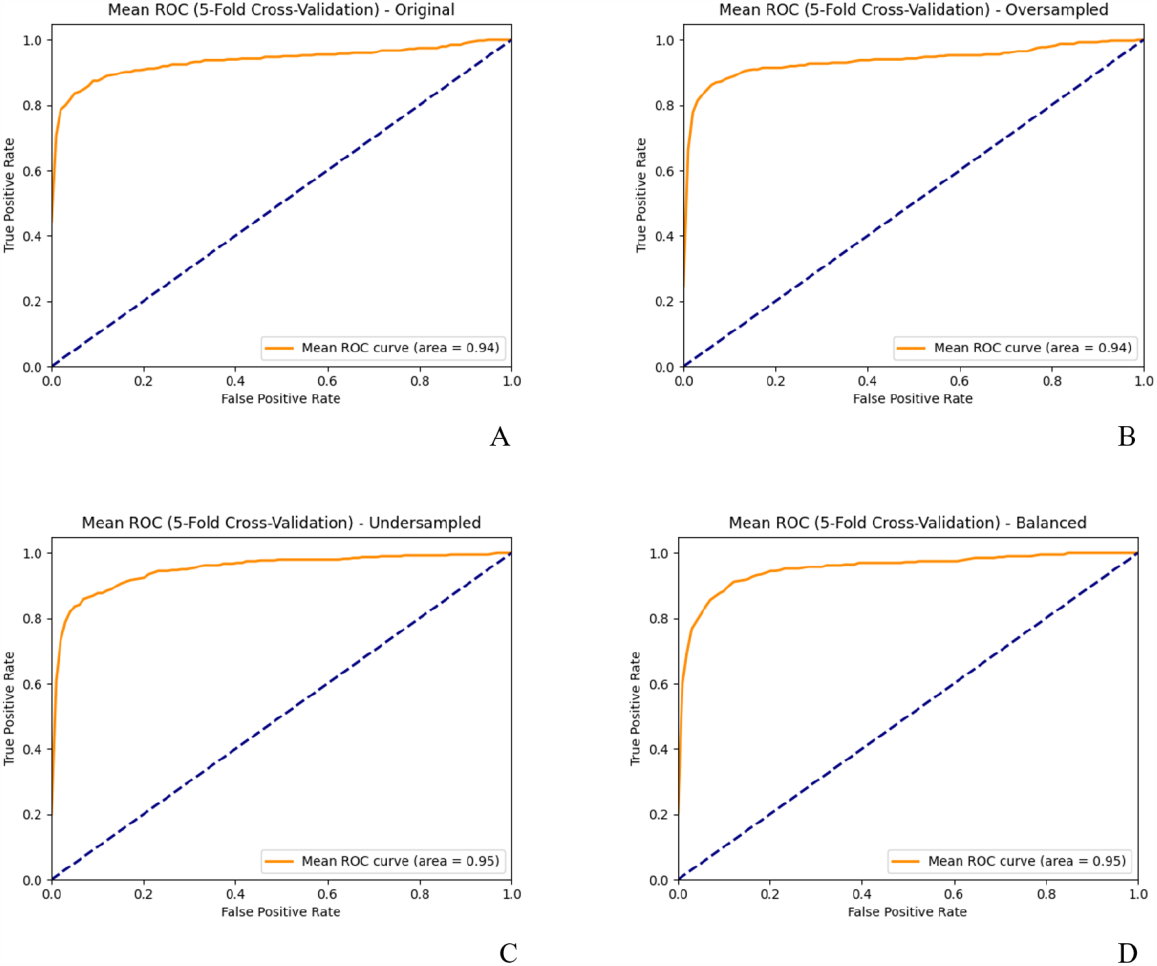
Mean AUROC across 5-fold cross-validation with the original dataset (A), oversampled dataset (B), undersampled dataset (C), and balanced dataset (D).

For visual representation, Figure 3 shows the mean values for the area under Receiver Operating Characteristic (AUROC) across 5-fold under different sampling methods. As clearly seen in the figure, the AUROC values for both original and oversampled datasets are 0.94, and the values for both undersampled and balanced datasets are 0.95.

### Leave-one-pathogen-out Validation

The LOPOV method was used to estimate how Vaxign-DL can be used to predict vaccine candidates for a novel pathogen. Performance for LOPOV showed good performance. Although the original and oversampled datasets exhibit identical mean AUROC values (Figure 4A, 4C), their performance diverges when considering individual pathogens. The original dataset not only outperforms the oversampled dataset in terms of mean AUPRC (Figure 4B, 4D) but also demonstrates superior pathogen-specific results. For instance, the performance of *Mycobacterium tuberculosis, Staphylococcus aureus, Helicobacter pylori*, and *Neisseria meningitidis* notably declines in the oversampled dataset.

**Figure 4.**
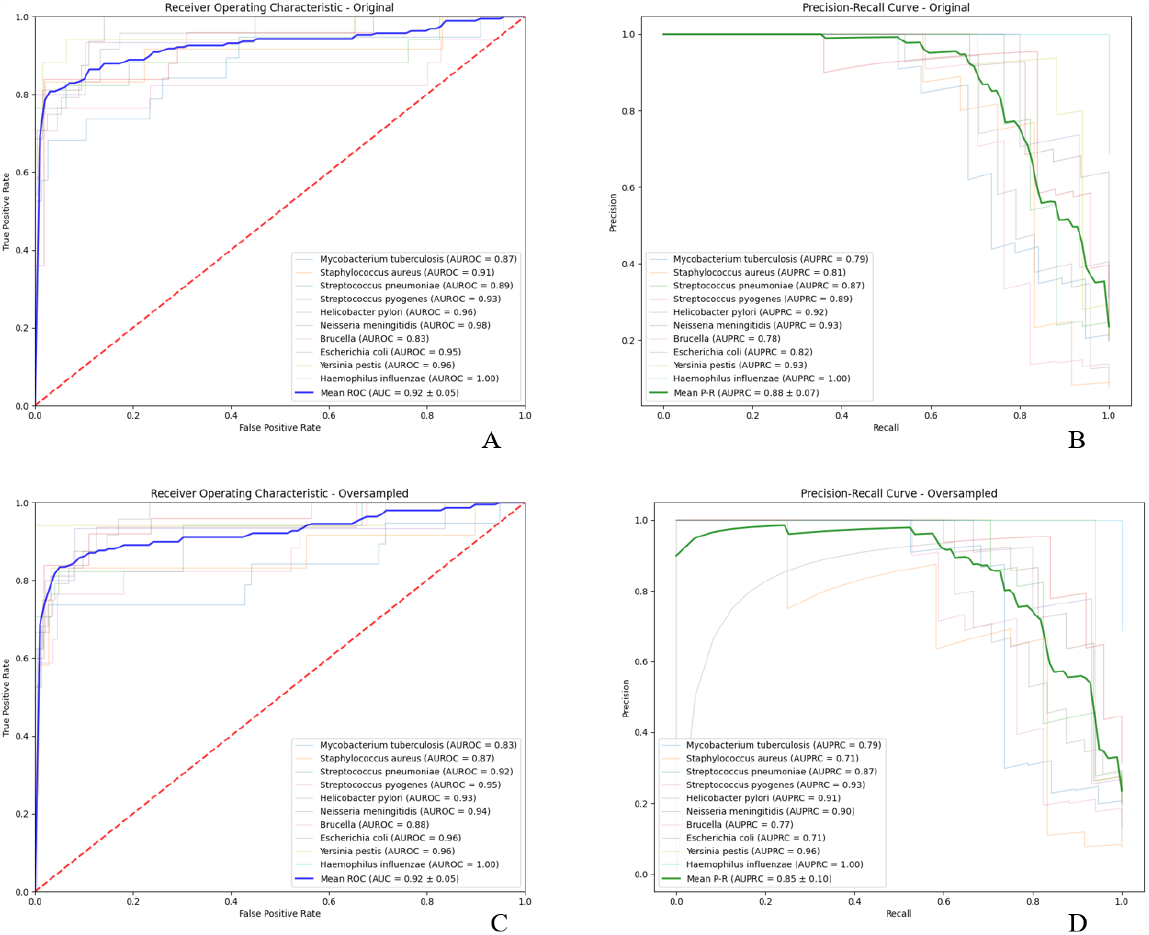
Receiver Operating Characteristic (A) and Precision-Recall Curve (B) with the original dataset. Receiver Operating Characteristic (C) and Precision-Recall Curve (D) with the oversampling method

### Benchmark different reverse vaccine design tools

We calculated the recall, precision, WF1, and MCC of existing RV tools to compare the performance of Vaxign-DL and Vaxi-DL (9) using the Vaxign-ML Benchmark dataset (8) (Table 2). Both Vaxign-DL (7) and Vaxign-ML tools exhibited significantly better performance than both Vaxi-DL and VaxiJen (11). This is caused by Vaxi-DL and VaxiJen having significantly worse precision. As we had utilized the same dataset as the previous Vaxign-ML paper (8), we included the performance of these prior RV tools. On this public dataset, Vaxign-DL exhibits comparable, but not superior performance to Vaxign-ML. Vaxign-DL currently is second on all four metrics we have measured.

**Table 2.** Assessment of Vaxign-DL, Vaxi-DL, Vaxign-ML, and other RV models using Vaxign-ML Benchmark Dataset.

|  | Recall | Precision | WF1 | MCC |
| --- | --- | --- | --- | --- |
| Vaxign-DL | 0.83 | 0.73 | 0.74 | 0.49 |
| Vaxi-DL | 0.92 | 0.67 | 0.70 | 0.46 |
| Vaxign-ML | 0.81 | 0.75 | 0.76 | 0.51 |
| Heinson–Bowman | 0.72 | 0.69 | 0.68 | 0.37 |
| VaxiJen | 0.69 | 0.68 | 0.66 | 0.32 |
| Vaxign | 0.32 | 0.79 | 0.56 | 0.27 |
| Antigenic | 0.50 | 0.52 | 0.49 | -0.02 |
| Epitope-based | 0.63 | 0.65 | 0.62 | 0.24 |

## Discussion

In this study, we developed the deep learning (DL)-based vaccine design program Vaxign-DL and demonstrated that Vaxign-DL has exhibited good performance in the prediction of bacterial protective antigens, showing the importance of integrating both biological and physicochemical features in deep learning (DL)-based reverse vaccinology (RV) prediction.

Compared to the usage of undersampled, oversampled, and balanced datasets, the use of the original dataset in our Vaxign-DL development has proven to be more effective, with the model demonstrating robustness and high scores performance metrics. The comparative analysis of different sampling methods has highlighted distinct advantages and disadvantages in the context of model performance. The undersampling method may discard critical data from the majority class. In contrast, oversampling does not significantly bring high-quality minority-class samples and carries the risk of overfitting.

Multiple differences exist between Vaxign-DL and Vaxi-DL (9). The training and the testing data appear quite different. Vaxi-DL uses the data in the VIOLIN Vaxgen (Vaccine-related Genes and Protective Antigens) database. However, not all the genes in Vaxgen are protective.

Instead, we used the VIOLIN Protegen protective antigen database data (10), in which all genes are protective.

In our future work, we will calculate and investigate additional features. Vaxi-DL used epitope prediction for prediction, which is not in our analysis. We will later explore how the inclusion of epitope prediction and other features may help our vaccine design performance in Vaxign-DL.

## Conclusion

Our work shows promise and replicability for applying Deep Learning for additional RV design. We are currently working on integrating additional features to improve the efficacy of prediction. We are also considering the use of DL methodology to include predictions if a protective antigen would be a virulence factor as a secondary predictor of vaccine safety.

## Author contributions

YZ: Neural network implementation and refinement, hyperparameter tuning, LOPOV validation, benchmark, and performance evaluation. AH: Training data generation, code refinement and debugging, hyperparameter exploration, model comparison, and mentoring of YZ in technical design and domain knowledge. JJ: Advising and consultation on neural network model generation. YH: Project design and management, model comparison, and data and result interpretation. YZ, AH, and YH participated in the first version of manuscript writing. All authors approved the paper publication.

## Acknowledgements

This project was funded by a NIH-NIAID grant U24AI171008. We also acknowledge the usage of the training data originally generated in the published Vaxign-ML study.

